# *Systems Biology* Inferring edge function in protein-protein interaction networks

**DOI:** 10.1101/321984

**Authors:** Daniel Esposito, Joseph Cursons, Melissa Davis

## Abstract

**Motivation:** Post-translational modifications (PTMs) regulate many key cellular processes. Numerous studies have linked the topology of protein-protein interaction (PPI) networks to many biological phenomena such as key regulatory processes and disease. However, these methods fail to give insight in the functional nature of these interactions. On the other hand, pathways are commonly used to gain biological insight into the function of PPIs in the context of cascading interactions, sacrificing the coverage of networks for rich functional annotations on each PPI. We present a machine learning approach that uses Gene Ontology, InterPro and Pfam annotations to infer the edge functions in PPI networks, allowing us to combine the high coverage of networks with the information richness of pathways.

**Results:** An ensemble method with a combination Logistic Regression and Random Forest classifiers trained on a high-quality set of annotated interactions, with a total of 18 unique labels, achieves high a average F1 score 0.88 despite not taking advantage of multi-label dependencies. When applied to the human interactome, our method confidently classifies 62% of interactions at a probability of 0.7 or higher.

**Availability:** Software and data are available at https://github.com/DavisLaboratory/pyPPI

**Supplementary information:** Supplementary data are available at *Bioinformatics* online.

## 1 Introduction

Following the genomic era, there has been significant effort to identify the functional characteristics of individual proteins. Many experimental based techniques have been developed to determine the function of proteins, however experimental methods are expensive and time consuming. As a response to this, and leveraging the large collections of sequence data that have emerged in the last two decades, numerous computational methods have been developed to infer the function of individual proteins. Proteins, however, do not function alone, and the normal functions of proteins are established through networks of interactions that form structural components, cascades of signalling, and regulatory complexes, among many other functions. This places protein-protein interactions (PPIs) at the heart of understanding molecular function.

This has inspired techniques using topological analysis of protein-protein interaction networks to infer protein function. Previous work has established that the structural organisation of the interactome can provide insight into the functional characteristics of proteins. For example, highly connected vertices, commonly referred to as network hubs, may be indicative of proteins with central biochemical roles (Barabasi & Oltvai, 2004). In one study, detailed topological analyses highlighted how the dynamics connectivity of the interactome can relate to disease pathology and progression (Ma, Gao, & Tan, 2014). Despite their large coverage, PPI networks lack the functionally rich information found in pathways, so we lose information regarding the functional nature and dynamics of individual molecular interactions. Conversely, pathways focus on much smaller groups of interacting proteins, annotating rich functional information such as the post-translational modifications (PTMs) occurring in each PPI at the cost of coverage As proteomic technologies continue to improve, we are uncovering the importance that PTMs have in shaping the function of cells.

Post-translational modifications typically involve the addition or removal of chemical groups, and are key regulators in many cellular processes (Bode & Dong, 2004). Moreover, studies have shown the importance of the interplay between PTMs have in the cross-talk and fine-tuning of pathway activity (Hunter, 2007). As research uncovers the intricacies of the effects on PTMs on cell function and behaviour, it is becoming more apparent that PTMs may represent something akin to a readable code allowing the fine-tuning of transcriptional regulation and protein activity (Winter, Erce, & Wilkins, 2014). Hence, protein-protein interactions from the point of view of PTMs is critical for understanding the biological function of cells and pathogenesis of disease. By combining the large coverage of PPI networks with the information richness of pathways, we can bring more biological insight into interactome-wide analyses.

Several algorithms have been proposed that investigate the type of protein-protein interactions based on various sources of data. Zhu et al. (2006) make use of X-ray crystallographic data to characterise interaction types into three classes: obligate, non-obligate and crystal packing. The drawback of this method is that it can only provide a low-level classification of the type of interaction. Silberberg et al. (2014) proposed a method to predict higher level PTM types based on experimental assay type in a machine learning approach. This method classifies protein-protein interactions into 10 high level types: cleavage, covalent binding, deacetylation, methylation, phosphorylation, dephosphorylation, ubiquitination, disulphide bonding, ribosylation and protein cleavage (at the time of publication, no implementation has been made available).

One limitation of the latter approach is that it does not make use of the rich information stored in functional and domain annotation databases such as the Gene Ontology (GO), InterPro and Pfam (Ashburner et al., 2000; Bateman et al., 2004; Zdobnov & Apweiler, 2001). There have been numerous successful approaches in using functional annotations in PPI inference. De Bodt et al. (2009) combined the maximum depth of the common ancestors of the GO annotations with sequence orthology and expression data to drive PPI prediction. Other methods have been employed that combine gene ontology terms into a semantic similarity score that uses the information content of ontology terms. Pesquita et al. (2008) provide a comprehensive review of various similarity measures used in PPI inference. Maetschke et al. (2012) have shown that exploiting the structural information contained with the GO directed acyclic graph (DAG) can provide superior PPI inference when compared to methods relying on semantic similarity based measures, or methods relying on annotations at the leaf nodes. The use of GO annotations has also been combined with domain and family annotations when performing PPI inference. Nan et al. (2004) and Lu et al. (2005) both provide comprehensive reviews of the various features used in PPI prediction tasks. Specifically, the use of the GO annotations was found to be among the most indicative features of a protein-protein interaction. Given the evidence presented by such studies, using functional annotations of proteins may prove to be excellent predictors of post-translational modifications in binary interactions. A second limitation is in the use of assay information as predictors, which are not as readily available as functional annotations.

Of the two prior approaches, neither method explores the use of multi-label classification algorithms. Attempting to predict the PTM types of a PPI introduces a significant challenge; two proteins may interact in several ways depending on the pathway and context meaning a PPI can have multiple PTM labels at once. This creates fuzzy decision boundaries which challenge conventional classification algorithms. A typical approach to multi-label classification is called Binary Relevance (BR), where a one-vs-rest classifier is trained for each label. Training an independent classifier for each label in this fashion has benefit of being simple to implement, and allows the analysis of the performance characteristics of each individual label. We also retain access to many advanced machine learning algorithms such as random forests (Breiman, 2001) and support vector machines that work well with single label data. The disadvantage of using BR is that each label is learned independently, effectively ignoring dependencies between labels that may exist if a dataset has significant label correlations. For example, seeing a phosphorylation label may be positively correlated with seeing an inhibition reaction.

In this paper, we present the first publicly available software package for the general prediction of post-translational modifications on protein interaction networks. The software package in the provided link is written in Python 3.6 and makes use of the Scikit-Learn machine learning library (Pedregosa et al., 2011).

## 2 Methods

*Data Collection:* The Kyoto Encyclopedia of Genes and Genomes (KEGG) is a knowledge base, organising different aspects of molecular biology into higher order information such as pathway maps (Kanehisa & Goto, 2000). We specifically make use of the databases relating to gene products, enzymes and pathways for *Homo sapiens*. The pathway database maps out molecular interactions and the type of interactions occurring between proteins. For this research, version 85.1 of KEGG (released 9/3/2018) has been used. KGG pathway maps are defined as a series of relations corresponding to molecular interactions. We filter these relations based on the interaction subtype, allowing only PPIs annotated to one or more of: ubiquitination, proteolytic-cleavage, methylation, prenylation, dissociation, inhibition, activation, state-change, glycosylation, sumoylation, deacetylation, acetylation, phosphorylation, dephosphorylation, binding/association, carboxylation, sulfation and myristoylation. The KEGG gene accessions were mapped to Uniprot Swissprot accessions using the KEGG’s LinkDB service. Samples for which TrEMBL only accessions were available were discarded to maintain a high quality training set.

We also mined labeled PPI samples from the Human Reference Protein Database (HPRD) (Keshava Prasad et al., 2008) PTM collection (release 9, 4/13/10). This was for several reasons: boosting the numbers of infrequent labels obtained from KEGG, obtaining more detailed reaction labels and to set aside an independent testing set for the labels already in the KEGG dataset and present in high numbers. Namely, the phosphorylation and dephosphorylation labels. The resulting PPIs were mapped to SwissProt using the mapping files provided by HPRD, which were then further mapped using UniProt to remove samples with obsolete, old or isoform variant accessions. Again, we filtered out PPIs which contained TrEMBL proteins to maintain a high quality testing and training set. Furthermore, labels having fewer than 5 samples were removed. Samples that were not labeled with phosphorylation and dephosphorylation were merged with the KEGG dataset to create the final multi-label training set.

To test the biological significance of model predictions, an unlabeled set of PPIs representing the human interactome from the Protein Interaction Analysis Platform (PINA, latest version released 21/05/2014) (Cowley et al., 2011), BioPlex network (Version 4a) (Huttlin et al., 2015)and InnateDB (version 5.4) (Lynn et al., 2008)were downloaded and parsed. These datasets were combined and processed, taking care to remove duplicate entires. Again, an additional mapping step was performed using the UniProt mapping service to remove non-human, obsolete and isoform variants.

*Feature extraction:* The UniProt database (Consortium, 2016) provides a central resource for cataloging information on proteins and protein sequences from several different sources. We have used SwissProt and TrEMBL (release 9/3/2018) to extract functional annotations from GO, InterPro and Pfam for all proteins. The UniProt mapping service has also been used on all datasets to map outdated and isoform variant accessions and to filter protein interactions containing non-human or obsolete accessions. We use annotations obtained from all three ontologies; Molecular Function, Biological Process and Cellular Component. As described in the following section, when using the inducer concept introduced by Maetschke et al. (2012) to construct feature vectors, only the ‘is a’ and ‘part of’ edge relationships of the GO DAG are used. Format version 1.2 released on 3/7/2018 has been used.

The features of a PPI are computed as the combined annotations of each protein, retaining duplicate annotations. When using textual features such as annotations, it is most common to represent these with a bag-of-words vector space model. To do this, we use binary encoding: a 0 if a term is not present for a PPI or a 1 if present from either protein. Additionally, we explore the use of ternary encoding. This encodes a 0 if a term is not present, 1 if it is present for one member and 2 if it is present for both.

To exploit additional information within the structure of the GO DAG, we use the concept of term inducers, which was shown to provide superior performance than using leaf annotations only or semantic similarity based methods (Maetschke et al., 2012). We use this under the assumption that inducing additional terms between two proteins will provide a classifier with richer functional information than simply taking the union of annotations for each protein. We specifically use the ‘up to lowest common ancestor’ (ULCA) term inducer, which finds the lowest common ancestors of two proteins within the GO DAG, and then includes all terms along the path from the leaf nodes to these ancestors. We compare this method to classifiers using only the leaf node annotations to explore its effectiveness.

*Classification:* In all experiments, bootstrapped 5-fold cross-validation repeated 3 times has been used. The folds were generated using multi-label iterative stratification (Sechidis, Tsoumakas, & Vlahavas, 2011). Using this algorithm, each training instance appears in the training and validation fold at least once. Within each fold, hyperparameters have been tuned using a randomised grid search (Bergstra & Bengio, 2012) on the training folds using a nested 3-fold cross-validation. During BR, a single classifier is trained for each label. Within each label, the class targets become binary indicator variables where a 1 indicates membership to that label or 0 otherwise. Using this method, each PPI is given a probability vector of label memberships. During classification, we use a probability threshold of 0.5 to determine label membership. We note that because of large class imbalances between present in our training dataset, scores such as accuracy and AUC may provide misleadingly optimistic results. Therefore, we rely on metrics such as the F1 score averaged across all folds, and then further averaged over each repeated iteration. We also measure multi-label performance using the ranking loss and the hamming loss functions applied to the binary predictions. Ranking loss is indicative of how well a classifier is predicting the positive labels of each sample, while the hamming loss represents the loss over positive and negative labels. Due to the nature of molecular biology and biases during PTM curation processes, it is inevitable that some labels will have significantly smaller sample sizes. For example, phosphorylation tends to be more commonly studied given that it is much easier to detect via mass spectroscopy and tends to play a central role in many biological processes (Hunter, 2007). This class imbalance presents difficulty for many classification algorithms, which will favour the majority class. Biased sampling approaches have been shown to provide little benefit for highly dimensional datasets (Blagus & Lusa, 2013). Alternative techniques that avoid sampling strategies are those based on cost-sensitive learning algorithms. By using a misclassification penalty inversely proportional to label frequency, we can place a higher penalty for misclassified instances belonging to the minority class.

*Interactome predictions:* We trained a classifier for each label on the combined training and testing dataset from KEGG and HPRD. Hyper-parameters are tuned by using 3-fold cross-validation with the best parameters being used to train the final classifiers using the whole dataset. The type of vector encoding was also included as a hyper-parameter. Unless stated otherwise, the features were computed from InterPro, Pfam and leaf node GO annotations. For each, PPI in the interactome, we predict a vector of probability estimates corresponding to each label in the training set.

To assess the strengths and weaknesses of our predictions over the interactome, we extracted a myristoylation interaction subnetwork containing all edges with a myristoylation probability of at least 0.5, and a sulfation interaction subnetwork containing all edges with a lower sulfation probability of at least 0.1 to demonstrate the topological effects of thresholding. Since our method relies on quality functional annotations to make meaningful predictions, annotation sparsity in uncharacterised or recently characterised proteins presents a potential issue.

## 3 Results

*Label distribution:* The final multi-label training set 26,716 unique multi-labelled samples The testing set from HPRD contained 1688 samples (1580 phosphorylation, 112 dephosphorylation). Unlike the training PPIs, the testing PPIs were not multi-label. The combined training set used during the interactome prediction contained 28,784 unique multi-labelled samples annotated to 18 labels, including those excluded during cross-validation (380 samples) to keep the training and testing sets disjoint. The distribution of labels amongst these protein interactions was found to be highly imbalanced with small handful of the labels making appearing in most of the interactions. This skewed distribution can be seen in table 1.

**Table 1.**
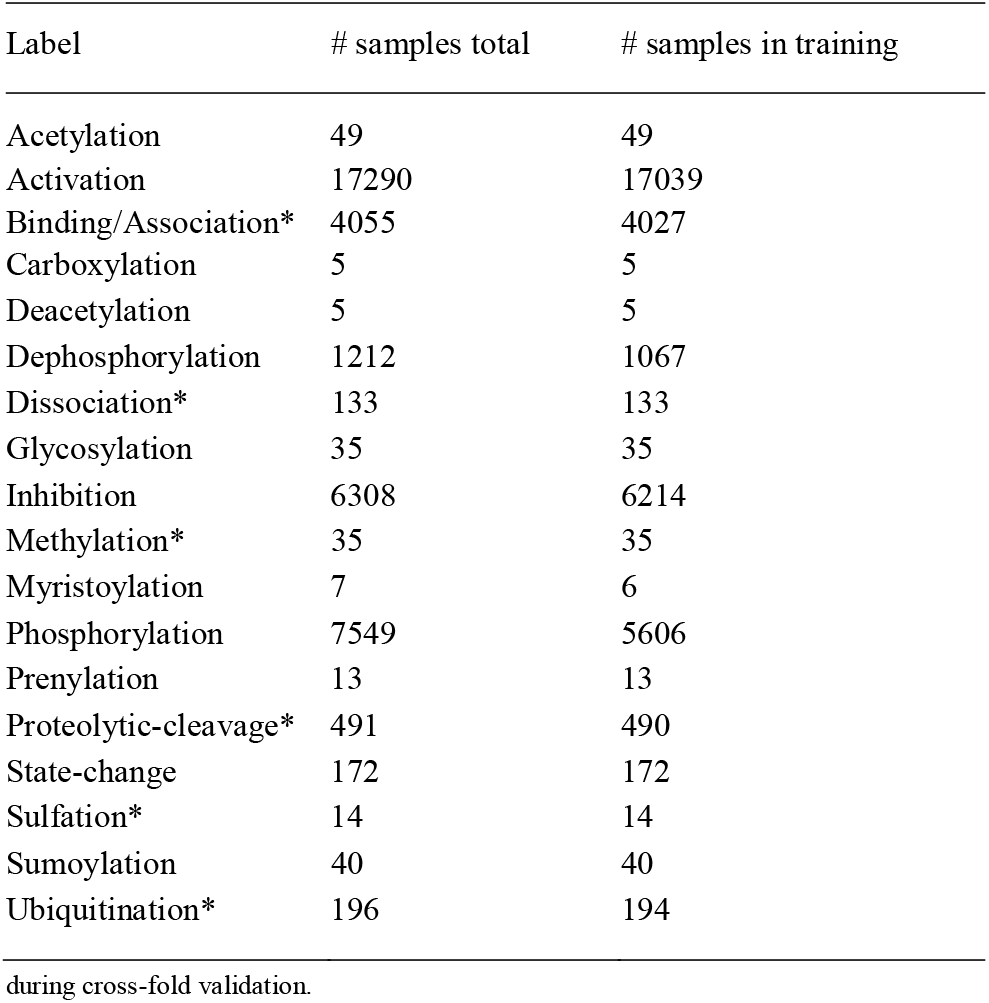
Label distribution in the full multi-label dataset and in the training set used during cross-fold validation.

| Label | # samples total | # samples in training |
| --- | --- | --- |
| Acetylation | 49 | 49 |
| Activation | 17290 | 17039 |
| Binding/Association* | 4055 | 4027 |
| Carboxylation | 5 | 5 |
| Deacetylation | 5 | 5 |
| Dephosphorylation | 1212 | 1067 |
| Dissociation* | 133 | 133 |
| Glycosylation | 35 | 35 |
| Inhibition | 6308 | 6214 |
| Methylation* | 35 | 35 |
| Myristoylation | 7 | 6 |
| Phosphorylation | 7549 | 5606 |
| Prenylation | 13 | 13 |
| Proteolytic-cleavage* | 491 | 490 |
| State-change | 172 | 172 |
| Sulfation* | 14 | 14 |
| Sumoylation | 40 | 40 |
| Ubiquitination* | 196 | 194 |

*data characteristics:*We computed the pairwise correlation between labels using the Spearman correlation coefficient to test for dependencies between labels. A heat map representing the strength of label correlations can be seen in figure 1a. We have used Jaccard’s similarity measure to assess how distinct the features annotated to each label are. For each label, the set of GO, InterPro and Pfam annotations from the PPIs annotated with the label were collected. Pairwise similarity scores between labels were computed using Jaccard’s measure over these respective annotation sets (figure 1b).

**Fig. 1.**
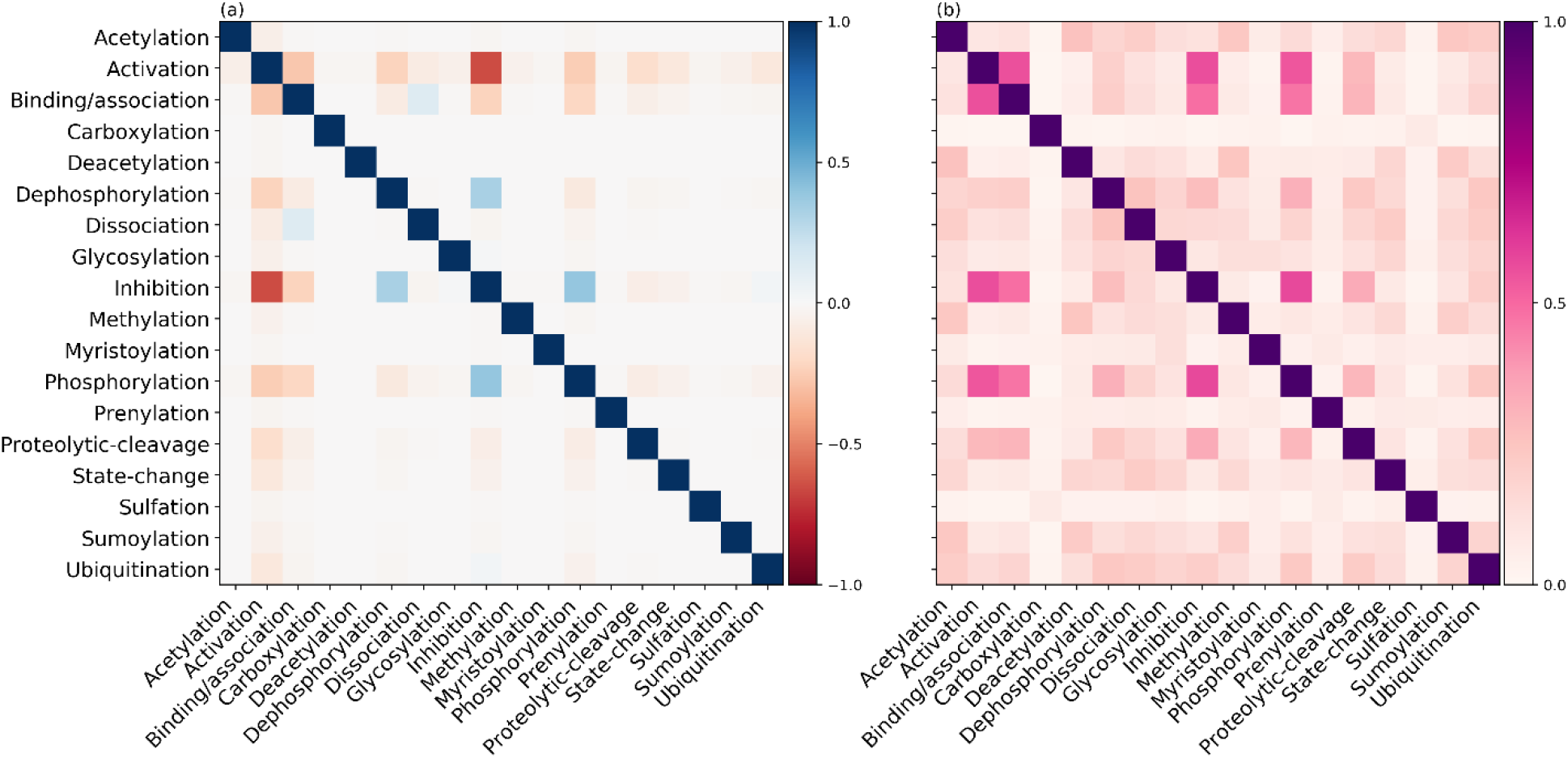
(a) The Spearman correlation between labels in the multi-label samples. (b) The Jaccard correlation coefficient between label pairs calculated from the proportion of shared features over all samples for each label.

*Cross-validation Performance:*For each label, we have trained a baseline classifier using a logistic regression classifier with binary feature encoding, where features are the annotations from the Gene Ontology, InterPro and Pfam. The F1 scores for each label are presented in table 2. Additional experiments have been performed with a single parameter varied relative to the baseline, and are also presented in table 2. Specifically, we have tested the effect of ternary encoding, the use of ULCA inducers and the use of random forests relative to the baseline. Additionally, each of the feature databases, InterPro, Pfam and have been trained over separately using the baseline configuration and these results are presented in supplementary table A.1. Testing performance for each experiment was computed for the labels phosphorylation and dephosphorylation over the heldout HPRD testing set. Other linear and nonlinear classifiers were also tested however generally, we found logistic regression and random forests to be the most efficient and best performing. When attempting to perform multi-label classification using classifiers such as structural SVMs and classifier chains, either worse or no significant performance gains were observed. Since they present significant computational burdens without any benefits, no further results on these methods will be presented.

**Table 2.** Cross-validated F1 scores for each label and overall multi-label performance. The baseline condition refers to a logistic regression classifier trained using ternary encoding with leaf-term GO, InterPro and Pfam features. Each column title refers to the condition modified for that experiment with respect to the baseline. Labels with ‘HPRD’ in parentheses represent the F1 performance over the HPRD testing set. Bold columns indicate overall highest performance determined by considering both the validation and testing scores.

| Label | Baseline | Ternary encoding | ULCA Inducer | Random Forest |
| --- | --- | --- | --- | --- |
| Acetylation | $0.78 \pm 0.015$ | $0.77 \pm 0.008$ | **$0.81 \pm 0.009$** | $0.77 \pm 0.029$ |
| Activation | $0.95 \pm 0.000$ | $0.94 \pm 0.000$ | $0.95 \pm 0.000$ | **$0.97 \pm 0.004$** |
| Binding/Association | $0.89 \pm 0.002$ | $0.87 \pm 0.000$ | $0.89 \pm 0.002$ | **$0.91 \pm 0.010$** |
| Carboxylation | **$1.00 \pm 0.000$** | $0.98 \pm 0.022$ | $1.00 \pm 0.000$ | $0.81 \pm 0.105$ |
| Deacetylation | $0.17 \pm 0.048$ | $0.18 \pm 0.049$ | $0.05 \pm 0.017$ | **$0.38 \pm 0.036$** |
| Dephosphorylation | **$0.98 \pm 0.001$** | $0.97 \pm 0.001$ | $0.97 \pm 0.001$ | $0.96 \pm 0.001$ |
| Dephosphorylation (HPRD) | $0.64 \pm 0.004$ | $0.63 \pm 0.005$ | $0.54 \pm 0.004$ | **$0.77 \pm 0.009$** |
| Dissociation | $0.84 \pm 0.010$ | $0.84 \pm 0.020$ | $0.84 \pm 0.012$ | **$0.87 \pm 0.008$** |
| Glycosylation | **$0.95 \pm 0.005$** | $0.95 \pm 0.006$ | $0.95 \pm 0.004$ | $0.87 \pm 0.023$ |
| Inhibition | $0.90 \pm 0.000$ | $0.89 \pm 0.000$ | $0.91 \pm 0.001$ | **$0.95 \pm 0.007$** |
| Methylation | $0.84 \pm 0.014$ | **$0.87 \pm 0.006$** | $0.80 \pm 0.013$ | $0.82 \pm 0.011$ |
| Myristoylation | **$1.00 \pm 0.000$** | $1.00 \pm 0.000$ | $0.91 \pm 0.059$ | $0.89 \pm 0.080$ |
| Phosphorylation | $0.92 \pm 0.001$ | $0.91 \pm 0.001$ | $0.93 \pm 0.001$ | **$0.95 \pm 0.008$** |
| Phosphorylation (HPRD) | $0.67 \pm 0.009$ | $0.64 \pm 0.010$ | $0.67 \pm 0.005$ | **$0.83 \pm 0.016$** |
| Prenylation | $0.97 \pm 0.033$ | $0.97 \pm 0.033$ | **$0.99 \pm 0.013$** | $0.75 \pm 0.062$ |
| Proteolytic-cleavage | **$0.83 \pm 0.002$** | $0.83 \pm 0.002$ | $0.83 \pm 0.007$ | $0.83 \pm 0.014$ |
| State-change | $0.89 \pm 0.002$ | **$0.92 \pm 0.001$** | $0.92 \pm 0.009$ | $0.85 \pm 0.004$ |
| Sulfation | $0.87 \pm 0.015$ | $0.84 \pm 0.015$ | $0.92 \pm 0.018$ | **$0.94 \pm 0.012$** |
| Sumoylation | $0.79 \pm 0.018$ | $0.69 \pm 0.014$ | $0.77 \pm 0.015$ | **$0.84 \pm 0.013$** |
| Ubiquitination | $0.88 \pm 0.009$ | $0.86 \pm 0.008$ | **$0.89 \pm 0.006$** | $0.87 \pm 0.028$ |
| F1 (averaged) | $0.86 \pm 0.002$ | $0.85 \pm 0.004$ | $0.85 \pm 0.004$ | $0.85 \pm 0.021$ |
| Ranking loss | $0.062 \pm 0.000$ | $0.069 \pm 0.001$ | $0.064 \pm 0.001$ | $0.046 \pm 0.005$ |
| Hamming loss | $0.011 \pm 0.000$ | $0.013 \pm 0.000$ | $0.010 \pm 0.000$ | $0.007 \pm 0.001$ |

*Interactome:* The interactome dataset contained 250,606 samples. We found that approximately 2.54% of the PPIs in the interactome appear in the training data. We note that PPI instances in the interactome suffer from additional feature vector sparsity due to annotations being present that were not seen during training, rendering them unusable. In the interactome dataset, InterPro and Pfam domains showed significant sparsity when compared to annotations from the Gene Ontology, with an average of 87% and 83% of features being usable. Conversely, Gene Ontology annotations showed much less sparsity with approximately 99% being useable. Alternatively stated, 17% of the PPI instances had less than half of their Pfam annotations being usable during prediction, 13% in their InterPro annotations and 0.1% in their GO annotations.

The final model used to classify the interactome was an ensemble classifier trained over all three annotation databases. For each label, we have used the results presented in table 2 as a guide to select the optimal classifier type. The full configuration is provided in supplementary table A.3. Vector encoding was included as an additional hyperparameter during randomised grid search. To give an indication of how well the final model was able to classify PTMs over the interactome, we measured the proportion of the interactome having maximum prediction probability for any label over a specific threshold. In practice, despite the apparent sparsity in the feature vectors, we were still able to confidently classify approximately 61% of the interactome at a probability of 0.7 or greater using the full model (figure 2). We investigated the distributions of predictions for each label using the same model over the individual annotations databases and provide this in table A.2 in the supplementary material.

**Fig. 2.**
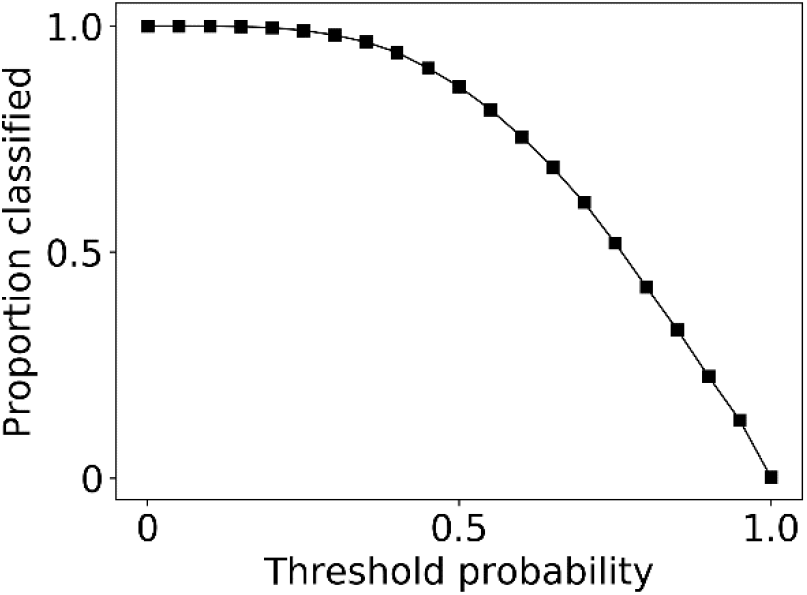
Relation between a probability threshold, t, and the proportion of the interactome classifiable at this threshold. A sample is counted as classifiable if the maximum probability observed across each label is greater than or equal to the given threshold.

Figure 3 shows the sub-networks induced for the sulfation and the myristoylation subnetwork. Red nodes indicate presence in the training dataset and red edges indicated the presence of that edge in the training data. Thicker edges indicate a higher label probability for sulfation and myristoylation respectively.

**Fig. 3.**
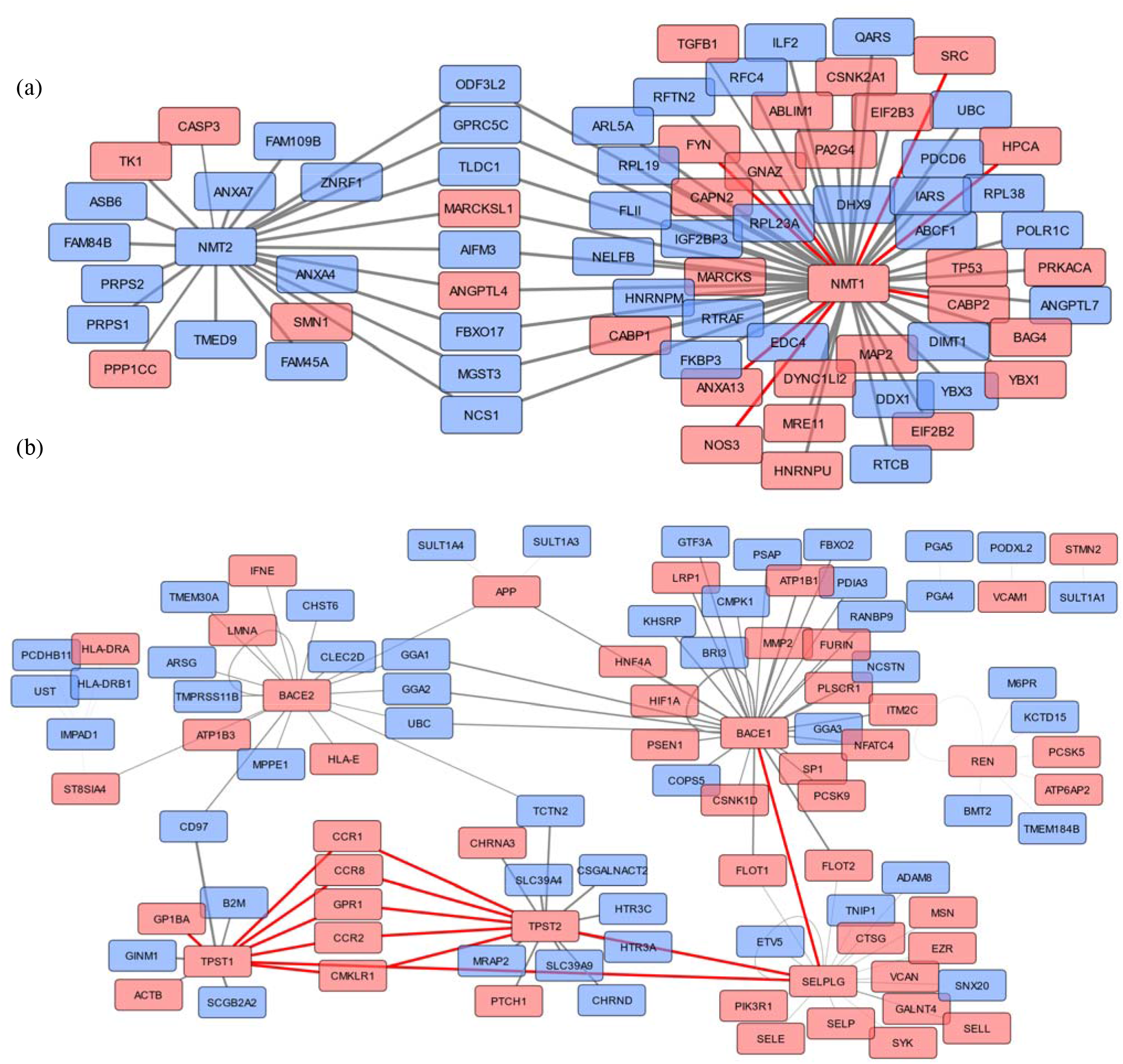
Post-translational modification predictions resulting from the application of the trained classifier on the interactome network. (Top and bottom) Nodes and edges highlighted in red appear in the training data. Grey edges represent newly classified post-translational modifications with a minimum probability of at least 0.5 for myristoyaltion and 0.1 for sulfation. Thicker edges indicate a higher classification probability (a) Top shows the induced myristoylation network showing an independently classified myristylation hub around NMT2. (b) Bottom shows the induced network for the label sulfation showing how the threshold probability can impact the connectedness of the subnetwork.

## 4 Discussion

In our modelling approach, we have used the binary relevance transformation turning each label into an independent binary classification problem. Despite binary relevance being a relatively naive approach to modelling multi-label datasets, we see excellent validation performance across each label, the exception being the label deacetylation. In terms of being able to identify multi-label samples, we see a very low ranking-loss.

In the test set, that the labels phosphorylation and dephosphorylation show a dip in F1 score, particularly the latter. We have attributed this to two main causes, the first being significant label correlations which create label dependencies. Since binary relevance ignores label correlations, we may lose additional information from the dependencies existing primarily between activation, phosphorylation, inhibition, dephosphorylation and binding/association as seen in figure 1a. The second reason is that phosphorylation and dephosphorylation (figure 1b) show higher annotation similarity with binding/association, activation and inhibition; the labels often confused with during testing predictions. The random forest classifier can more effectively learn these labels given these apparent non-linearities in the features.

When fitting the baseline model on Pfam annotations only (supplementary table A.1), labels such as sulfation, dissociation and methylation have significantly worse performance. We found that the two proteins O60507/TPST1 and O60704/TPST2 participating in most sulfation interactions had no Pfam annotations ex-plaining why the F1 scores for the partial model fit on Pfam annotations were so low. The label methylation showed similarities to sulfation in that the protein Q99873/PRMT common to a moderate amount of interactions had no Pfam annotations. For the label dissociation, the central proteins participating in most interactions had only one Pfam domain making it highly likely that the classifier was over-fitting to this feature. Using Pfam presents additional complexities since it generally provides a less diverse range of annotations, hence a combination approach by using all feature databases may be superior.

The goal of using the ULCA inducer was to include additional semantic and structural information from the GO-DAG into the feature vector. In table 2 we see a similar scenario when comparing the effect the ULCA inducer; most labels seem to show negligible benefits, with the exception of a select handful which may help in providing greater diversity in the features classified on, offsetting the previous issues of low protein diversity and low sample sizes. The use of inducers adds significant computational cost to both feature computation and classifier training, without providing any significant benefit to a random forest classifier trained on leaf terms. An alternate method is to use GO Slim variants of the ontology, however related studies have shown GO Slims to be inferior in prediction related tasks due to loss of information (Maetschke et al., 2012).

When using various partial and full models to make predictions over the interactome, we see large discrepancies in the distribution of predicted labels starred in table 1. For example, using a partial model fit only on Pfam we see very large number of predictions for sulfation, methylation, proteolytic-cleavage and dissociation. A likely cause for this difference is that, on average the density of Pfam feature vectors for PPIs in the interactome is significantly lower due to annotation sparsity.

Consequently, what we end up with is an inflation of unreliable predictions. This is where using a model based on all three annotation databases can help to balance out predictions and providing a richer source of information when Pfam, and to a lesser extent InterPro, annotations are less reliable. We also note that some labels, most notably binding/association, ubiquitination and proteolytic-cleavage have much higher predicted proportions (supplementary table A.2) than in the training data. This may represent an artefact of the curation process, which tends to focus on more common PTMs that are easier to detect (Mann & Jensen, 2003).

As illustrative examples, we have taken the sub-networks corresponding to myristoylation and sulfation to highlight the strengths and weaknesses of our classification method. Figure 3a shows the predicted myristoylation sub-network using a threshold of 0.5. The red edges and nodes represent those found in the training data. At this threshold, we have predicted the majority of edges and additionally, predicted the presence of a secondary module around the hub NMT2 not seen in the training data. Altering the probability threshold was found to significantly impact the network topology as can be observed in figure 3b. For example, setting the probability too low can result in module fragmentation and the identification of orphaned interaction pairs. Hence, we propose that when using our method, domain information may be useful to guide the selection on an appropriate threshold when inducing networks for analysis.

### Funding

This work has been supported by The University Of Melbourne and The Walter and Eliza Hall Institute of Medical Research.

*Conflict of Interest:* none declared.

